# Protein language modeling and supervised machine learning reveal the functional landscape of antimicrobial resistance genes in African metagenomes

**DOI:** 10.64898/2026.01.24.701459

**Authors:** Martin N. Muigano, Grace W. Kamau

**Affiliations:** Bio One Scientific, Nairobi, Kenya

**Author notes:** Corresponding author’s.

**Keywords:** Antimicrobial resistance, metagenomics, Africa, protein language models, resistome, machine learning, functional embedding

## Abstract

In this work, we analyzed 184 metagenomes from sub-Saharan Africa to characterize the functional landscape of ARGs using a combination of homology-based annotation and protein language model embeddings. We obtained 5,066 high-confidence ARG protein sequences from the African metagenomes, which we compared with 6,052 reference ARGs from CARD using embeddings generated with the ESM-2 protein language model. Additionally, we used a random forest classifier to determine the role of amino acid sequence and physiochemical features in discriminating ARGs and non-ARG sequences. The curated dataset revealed a predominance of ESKAPE pathogens in the resistome. β-lactam resistance was the most prevalent functional class, accounting for 4,046 ARG assignments (79.87%). At the country level, Burkina Faso, Malawi, and Benin exhibited the highest ARG hits per sample, thus demonstrating geographic heterogeneity in ARG burden. Protein language modeling demonstrated that African ARG sequences largely occupied the same functional subspace as globally curated CARD proteins. Supervised machine learning based on protein compositional and physicochemical features achieved high discriminatory performance between ARGs and non-ARGs (accuracy = 0.94, ROC-AUC = 0.99). Feature importance analysis identified amino acid composition, protein length, molecular weight, and isoelectric point as key discriminators, with statistically significant differences between ARG and non-ARG proteins (Mann-Whitney U test, p < 0.001). These results suggest that ARGs in African metagenomes are shaped primarily by ecological filtering and antibiotic selection pressure rather than by the emergence of novel resistance functions. This work provides a functional baseline for AMR surveillance in Africa and highlights the value of protein language models for resistome-scale analyses.

## 1. Introduction

Antimicrobial resistance (AMR) has emerged as one of the most formidable challenges facing public health systems today. AMR occurs when microorganism develop adaptation mechanisms to survive the presence of antibiotics that would naturally be effective against them (Dadgostar, 2019). This creates complex challenges for antimicrobial treatment. To put this into context, the steady erosion of antibiotic efficacy now threatens not only routine clinical care but also the economic and public health gains achieved over decades, with rising morbidity, mortality, and healthcare costs reported worldwide (Naylor et al., 2018). Meanwhile, the burden of AMR is particularly pronounced in sub-Saharan Africa. In 2019, an estimated 1.05 million deaths in Africa were attributed to bacterial AMR (Sartorius et al., 2024). The burden is made worse by increased costs to the already strained healthcare systems and prolonged hospitalizations (Gulumbe et al., 2022). In this region, close human–animal–environment interactions accelerate the spread of antimicrobial resistance genes (ARGs). Limited resource availability also hinders surveillance infrastructure (Muigano et al., 2026). In such settings, resistance does not arise in isolation but is rather embedded within complex microbial communities spanning human microbiomes, livestock, wildlife, soils, aquatic systems, and wastewater environments.

Advances in metagenomic sequencing have made it possible to determine resistance patterns in complex microbial communities. Over the years, function- and sequence-based analyses of metagenomic datasets have led to the discovery of numerous novel genes and metabolic functions (Momper et al., 2023; Pehrsson et al., 2013; Prayogo et al., 2020). These efforts have reshaped our understanding of microbial community structure and functional capacity. Within this framework, a major objective of metagenomic studies is the identification and annotation of protein-coding genes, including ARGs. However, most ARG detection pipelines remain firmly rooted in homology-based strategies. These typically rely on sequence similarity searches against curated reference databases such as the Comprehensive Antibiotic Resistance Database (CARD) or ResFinder (Manzoor et al., 2025). Tools like DIAMOND that are based on best-hit alignments against known resistance determinants have proven effective in identifying conserved ARGs. However, while we cannot rule out their utility in well-characterized systems, the limitations of such approaches are increasingly apparent. Homology-based methods struggle to detect divergent or novel ARGs, particularly when sequence identity drops below conventional thresholds (Berglund et al., 2019). This constraint is especially problematic in metagenomic reads whose short fragment lengths may be harder to annotate. Additionally, these methods may be limited in underrepresented regions such as Africa where microbial populations may be shaped by distinct evolutionary histories and selective pressures.

Against this backdrop, alternative strategies that move beyond strict sequence similarity have gained traction. Protein-intrinsic features such as physicochemical properties, amino acid composition, secondary structure content, and stability indices carry functional information that is relevant to AMR prediction (Bonifacio-Velez de Villa et al., 2025). Importantly, these features are often conserved even when primary sequence similarity erodes. Meanwhile, machine learning tools have been shown to be promising in exploiting such features to distinguish ARGs from non-ARG proteins with high accuracy (Rannon et al., 2025). (Rannon et al., 2025). Additionally, recent advances in protein language models (PLMs) have allowed the embedding of protein sequences into high-dimensional vector spaces that capture structural and functional properties without the need for explicit alignment or annotation. Such embeddings offer a useful framework for comparing proteins at scale and for visualizing functional landscapes beyond simple sequence similarity. PLMs can be used to examine whether ARGs from specific regions, such as Africa, are functionally distinct from globally curated resistance genes or whether they reflect conserved resistance solutions shaped by shared selective pressures. In this study, we combined metagenomic ARG detection with protein language model embeddings and machine learning to investigate the functional landscape of ARGs recovered from African metagenomes. We asked three primary questions: (i) How are resistance functions structured within African metagenomes? (ii) Do African metagenome-derived ARGs occupy distinct regions of functional protein space relative to the global CARD resistome? (iii) Can amino acid and physciochemical features reliably discriminate ARG versus non-ARGs in African resistome? By integrating homology-based annotation, embedding-based analysis, and machine learning, we provide a systematic assessment of the African resistome and its relationship to global resistance diversity.

## 2. Materials and Methods

### 2.1 Data acquisition and processing

In December 2025, we searched for whole genome sequence (WGS) metagenomes of African origins from the NCBA SRA database using a combination of key words: (“metagenomic” OR “metagenome” OR “shotgun sequencing”) AND (“whole genome sequencing” OR “WGS”) AND (“Country” OR “Sub-Saharan Africa”). The “Country” keyword was varied with specific countries in sub-Saharan Africa. From each bioproject, at least 2 WGS metagenomes were randomly selected and their raw reads downloaded from ENA using weget bash scripts. A total of 184 shotgun metagenomes were compiled from publicly available datasets representing diverse environments across sub-Saharan Africa, including human-associated, animal-associated, and environmental samples (e.g., soil, water, wastewater). Sampling locations spanned multiple countries and ecological contexts. This allowed us to obtain broad representation of African resistomes. Metadata including sample origin, environment type, and geographic location were curated from original study records.

### 2.2. Identification of antimicrobial resistance genes

Raw paired-end metagenomic sequencing reads were processed using a custom bash script. To ensure alignment-free resistome screening, FASTQ files for each sample were downloaded directly and processed without assembly or gene prediction. Instead, antimicrobial resistance genes (ARGs) were identified using a read-based translated alignment strategy implemented in DIAMOND (v2.x) (Buchfink et al., 2021). Specifically, raw nucleotide reads were queried against the Comprehensive Antibiotic Resistance Database (CARD) protein reference (Alcock et al., 2020) using DIAMOND BLASTx. This allowed the translation of the reads in all six reading frames and subsequent direct alignment to known resistance proteins.

To ensure high-confidence ARG detection, we filtered the alignments using stringent criteria by imposing a minimum amino acid identity of 80% and an E-value threshold of 1 × 10 ¹. For each retained hit, sample identifiers, read identifiers, matched CARD protein accessions (ARO), percent identity, E-value, and bitscore were recorded. Samples that did not yield any ARG hits under these criteria were excluded from downstream analyses. Filtered ARG hits from all samples were aggregated into a unified table that contained the read-level evidence for the presence of CARD resistance determinants across African metagenomes. This strategy ensured high-confidence ARG detection, minimization of false positives, and strong functional confidence.

### 2.3. Construction of the African ARG reference subset

To enable functional embedding analyses, we constructed a non-redundant African ARG reference subset consisting exclusively of CARD protein sequences that were detected in at least one African metagenome. This process yielded 5,066 high-confidence ARG protein sequences derived from African metagenomes. Each sequence header retained information on the associated CARD ARO accession, resistance gene name, and predicted host taxonomy when available. It is important to note that this reference does not represent *de novo* predicted proteins but rather known resistance proteins from CARD with empirical read-level support in African samples. This strategy allows assessment of the functional space occupied by African resistomes while maintaining direct comparability with the global CARD database. For comparison, a global reference resistome was constructed using the CARD protein homolog model dataset, consisting of 6,052 curated ARG protein sequences. This dataset represents a broad collection of experimentally validated and well-characterized resistance genes spanning multiple resistance mechanisms and drug classes.

### 2.4. Protein language model embeddings

We generated protein language model embeddings using the ESM-2 transformer model (ESM-2, 33-layer, 650M parameter model). The Evolutionary Scale Modeling framework (ESM-2) encodes protein sequences into dimensional vectors to capture sequence, structural, and functional constraints (Lin et al., 2023). Embeddings were computed for both the African ARG reference subset and the complete CARD protein collection. Each protein sequence (African ARGs and CARD reference ARGs) was embedded independently, producing a 640-dimensional vector representation per sequence. Protein sequences were converted into model tokens using the ESM alphabet batch converter and passed through the network without gradient computation. Mean pooling across residue embeddings was used to generate a fixed-length embedding for each protein. These embeddings were subsequently used for dimensionality reduction, functional landscape visualization, density bias analysis, and proximity assessment between African and global resistance proteins. Embeddings were generated in batches and saved as NumPy arrays for downstream analysis. In total, embeddings were successfully generated for 5,066 African ARGs and 6,052 CARD ARGs.

### 2.5 Dimensionality reduction and Functional density analysis

To visualize the functional landscape of the ARGs generated in the previous step, we projected the embeddings into two-dimensional space using Uniform Manifold Approximation and Projection (UMAP) (McInnes et al., 2018). UMAP was applied to the combined African and CARD embedding matrix using a fixed random seed for reproducibility. Visualizations were used to assess global overlap, clustering patterns, and potential separation between African and global ARGs. To quantify relative occupancy of functional space, local density estimates were computed in embedding space for African and CARD ARGs separately using k-nearest neighbor density estimation. For each point in the combined embedding space, a log density ratio was calculated as log(Africa density / CARD density). This allowed us to identify regions enriched for African or CARD ARGs. This approach enabled the assessment of whether African ARGs preferentially occupy distinct or biased regions of functional protein space.

### 2.6. Functional annotation and enrichment analysis

Resistance mechanisms and drug class annotations were assigned to African ARGs by mapping ARO accessions to CARD ontology terms. Drug class annotations containing multiple entries were split to account for multi-drug resistance genes. Functional distributions were summarized using frequency and proportion analyses. Comparisons between African and CARD ARGs were conducted to identify similarities and differences in resistance mechanism composition. Within African ARGs, joint distributions of resistance mechanism and drug class were examined to assess internal functional structure.

### 2.7. Supervised Machine Learning

We employed a supervised machine-learning approach to distinguish antimicrobial resistance (ARG) proteins from non-ARG proteins based on physicochemical characteristics and amino acid sequence–derived features. Supervision was achieved by training the model on proteins with known ARG status: CARD-detected ARGs from the African metagenomes (positive class) and curated non-ARG microbial proteins from Uniprot (negative class). A Random Forest (RF) classifier was selected due to its robustness to correlated features, ability to model non-linear relationships, and interpretability via feature importance measures. Random Forests also perform well on tabular biological data without strong parametric assumptions.

For training data construction, we used two sets of labeled training sets:

- Positive class: In this class, protein sequences were detected from uniformly processed African metagenomes using DIAMOND alignment against the Comprehensive Antibiotic Resistance Database (CARD). A total of 5,066 sequences formed this positive class.
- Negative class: The negative training set was derived from UniProtKB/Swiss-Prot and consisted of reviewed bacterial proteins excluding entries annotated with antibiotic resistance–related keywords or resistance-associated protein names. These sequences represent typical microbial proteins unlikely to be involved in resistance, reducing label ambiguity. This approach minimized contamination of the negative class while preserving broad functional diversity. A sample of 5,070 sequences was obtained randomly from the UniProtKB

For each protein sequence, we obtained 25 physiochemical and compositional features. These were computed using Biopython and custom bash scripts. These features were in two major categories:

- Global protein features: These included the sequence lengths as well as the fractions of polar, non-polar, positively charged, and negatively charged residues.
- Amino-acid composition: We also obtained features associated with the fractional abundance of each amino acid including alanine (AA_A), cysteine (AA_C), aspartate (AA_D), glycine (AA_G), arginine (AA_R), glutamate (AA_E), and tryptophan (AA_W) among others. These features captured biochemical constraints, charge distribution, and compositional biases associated with resistance proteins.

The random forest classifier was trained using a stratified 5-fold cross-validation approach. The classifier was implemented using the scikit-learn library and trained with default hyperparameters. Random Forests were selected due to their robustness to feature scaling, ability to model nonlinear decision boundaries, and inherent estimation of feature importance. For each fold, the trained model produced probabilistic predictions for the held-out test set, allowing performance to be assessed across multiple decision thresholds.

Model performance was quantified using accuracy, precision, recall, F1-score, receiver operating characteristic (ROC) curves, and area under the ROC curve (AUC). Precision-recall (PR) curves were also generated to evaluate classifier behavior under varying probability thresholds. Mean ROC and PR curves were computed by interpolating fold-wise results. The trained model was then fitted on the full balanced dataset and serialized for downstream application. This final model was used to predict ARG probabilities for proteins derived from independent African metagenomes and metagenome-assembled genomes (MAGs) for large-scale, alignment-free screening for potential antimicrobial resistance genes. The performance metrics were calculated as follows:

Accuracy: overall proportion of correctly classified instances

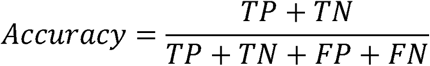

Precision: the proportion of predicted positives that are true positives

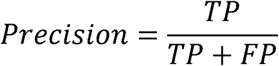

Recall (sensitivity): the proportion of correctly identified true positives

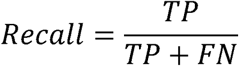

Where; TP = true positives, TN = true negatives, FP = false positives, and FN = false negatives

### 2.8. Statistical analysis

Statistical analyses were performed using Python (v3.9) with the SciPy and scikit-learn libraries. Differences in protein feature distributions between ARGs and non-ARGs were assessed using the non-parametric Mann–Whitney U test. This was necessitated by the non-normal distribution of features analyzed. All tests were two-sided, and statistical significance was defined as p < 0.001. Machine-learning model performance was evaluated using stratified cross-validation, and classification metrics included accuracy, precision, recall, F1-score, and area under the receiver operating characteristic curve (ROC-AUC). Feature importance rankings were derived from the trained model and interpreted in conjunction with univariate statistical tests to support biological relevance.

## 3. Results

### 3.1 Composition of the African resistome

We processed 184 metagenomes from various sub-Saharan countries including South Africa (n = 21), Ethiopia (n = 16), Zimbabwe (n = 12), Tanzania (n = 12), Uganda (n = 10), Kenya (n = 10), Ghana (n = 10), Burkina Faso (n = 8), Botswana (n = 8) Niger (n = 8), Mali (n = 6), Cameroon (n = 5), Gabon (n = 5), Benin (n = 5), Chad (n = 4), Mozambique (n = 4), Nigeria (n = 4), Republic of Congo (n = 4), Madagascar (n = 4), Democratic Republic of Congo (n = 3), Cabo Verde (n = 3), Seychelles (n = 3), Cote d’Ivoire (n = 3), and Malawi (n = 3). Other samples came from Gambia, Liberia, Central African Republic, Togo, Namibia, and Senegal (n = 2 in each country) as well as a single sample from Rwanda.

The African AMR hotspot analysis revealed substantial geographic heterogeneity in both ARG burden and diversity after normalization for sampling effort (Figure 1). Burkina Faso exhibited the highest normalized ARG intensity with an average of 221,396 ARG hits per sample. This despite being represented by only eight metagenomes. This suggests a concentrated resistance signal rather than an effect of oversampling. Similarly, Malawi (184,318 hits per sample), Benin (154,690 hits per sample), Seychelles (140,827 hits per sample), and Mali (136,377 hits per sample) ranked among the most intense AMR hotspots, each showing disproportionately high ARG loads relative to their sample counts. Several countries with larger numbers of analyzed metagenomes, including Zimbabwe (12 samples; 120,467 hits per sample), Ghana (10 samples; 108,611 hits per sample), Kenya (10 samples; 95,107 hits per sample), and Tanzania (12 samples; 93,546 hits per sample), also demonstrated elevated AMR intensity. This suggests sustained and widespread resistance signals across multiple ecological contexts. In contrast, countries such as South Africa (21 samples; 45,582 hits per sample), Ethiopia (16 samples; 33,451 hits per sample), and Nigeria (4 samples; 17,305 hits per sample) exhibited comparatively lower normalized intensities despite substantial sequencing representation. Collectively, the findings indicate that AMR burden across Africa is not uniformly distributed and that high-intensity resistance hotspots.

**Figure 1:**
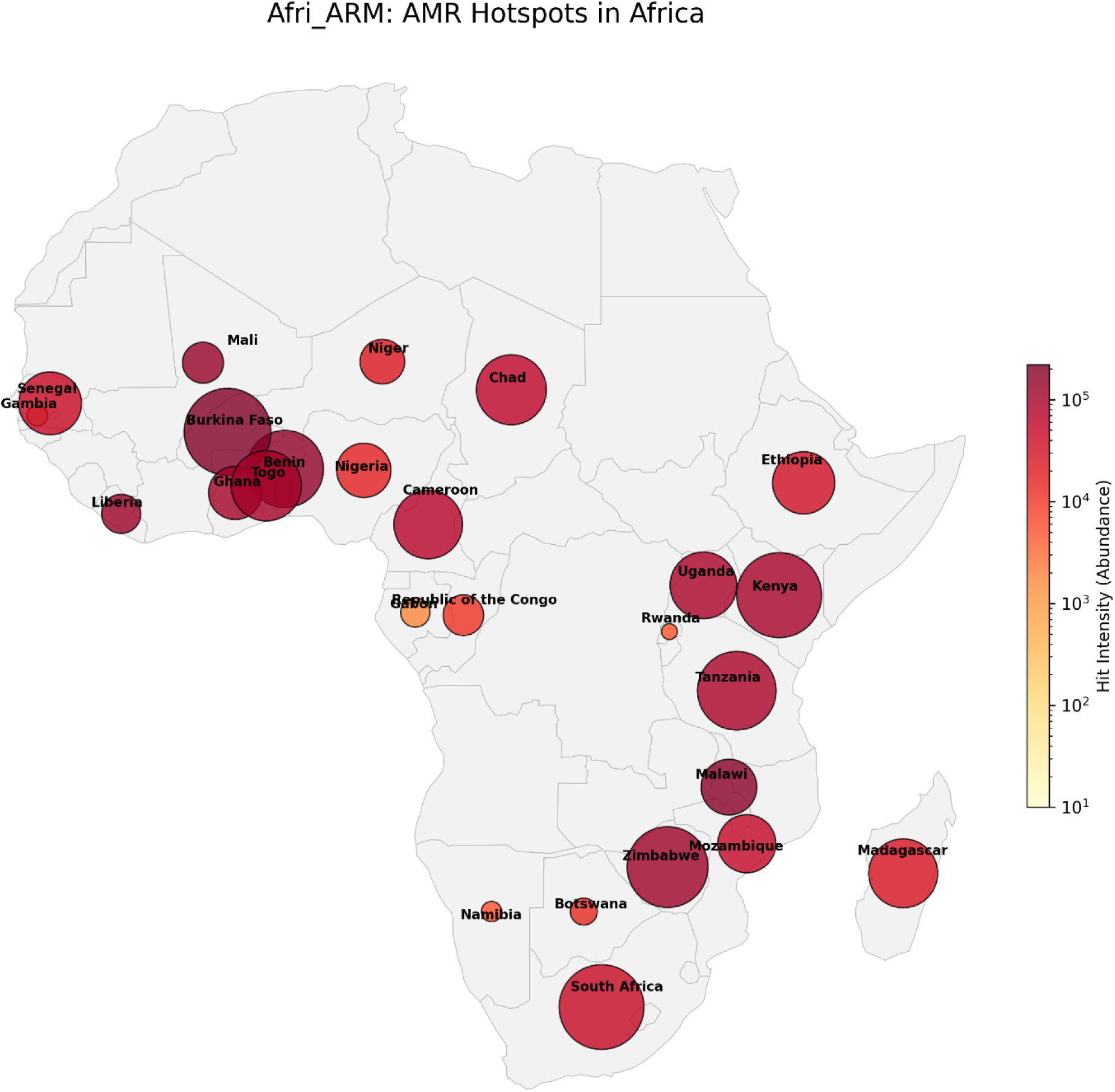
Geographic distribution of antimicrobial resistance (AMR) hotspots across Africa. Each circle represents a country with available metagenomic data included in the Afri-ARM analysis. Circle size corresponds to the number of metagenomic samples analyzed per country, while color intensity reflects the normalized ARG detection intensity, calculated as the total number of CARD-aligned ARG hits divided by the number of samples from that country. Darker colors indicate higher ARG hit intensity per sample.

### 3.2. Taxonomic drivers of antimicrobial resistance

Taxonomic attribution of ARGs indicated that resistance determinants were predominantly associated with clinically relevant opportunistic and enteric bacteria. *Pseudomonas aeruginosa* was the most frequent contributor (n = 712), followed by *Acinetobacter baumannii* (n = 570), *Escherichia coli* (n = 559), and *Klebsiella pneumoniae* (n = 529). Members of the *Enterobacter cloacae* complex (n = 190) and *Citrobacter freundii* (n = 121) further contributed to the observed resistance landscape. Notably, a substantial number of ARGs were assigned to unclassified or uncultured bacteria (n = 192 combined), which could be indicative of poorly characterized microbial taxa contributing to the African resistome. The prominence of *Acinetobacter*, *Pseudomonas*, and Enterobacteriaceae species suggests that there could be substantial overlap between environmental, commensal, and clinically relevant reservoirs of antimicrobial resistance across the continent. These microrganisms are frequently implicated in multidrug-resistant infections.

Figure 2 shows the mapping of the pathogens to antibiotic resistance drug classes. Antibiotic resistance annotations revealed a resistome strongly dominated by β-lactam resistance, which accounted for 4,046 ARG assignments, equivalent to 79.87% of all drug classes. This was followed at a much lower frequency by resistance to aminoglycosides (n = 193), fluoroquinolones (n = 137), and peptide antibiotics (n = 133). Additional resistance classes like lincosamides (n = 85), tetracyclines (n = 77), glycopeptides (n = 74), and diaminopyrimidines (n = 62) were present but contributed comparatively minor fractions of the overall resistome. Resistance to macrolides and phenicols (each n = 50), as well as to biocides/antiseptics (n = 39) and phosphonic acids (n = 33) was also detected at low abundance. Rare drug classes such as rifamycin, nitroimidazole, and aminocoumarin were also present. This distribution points to a skewness toward resistance mechanisms targeting β-lactam antibiotics. It could also be an indication of extensive and sustained selective pressure across African human, animal, and environmental settings for this drug class.

**Figure 2.**
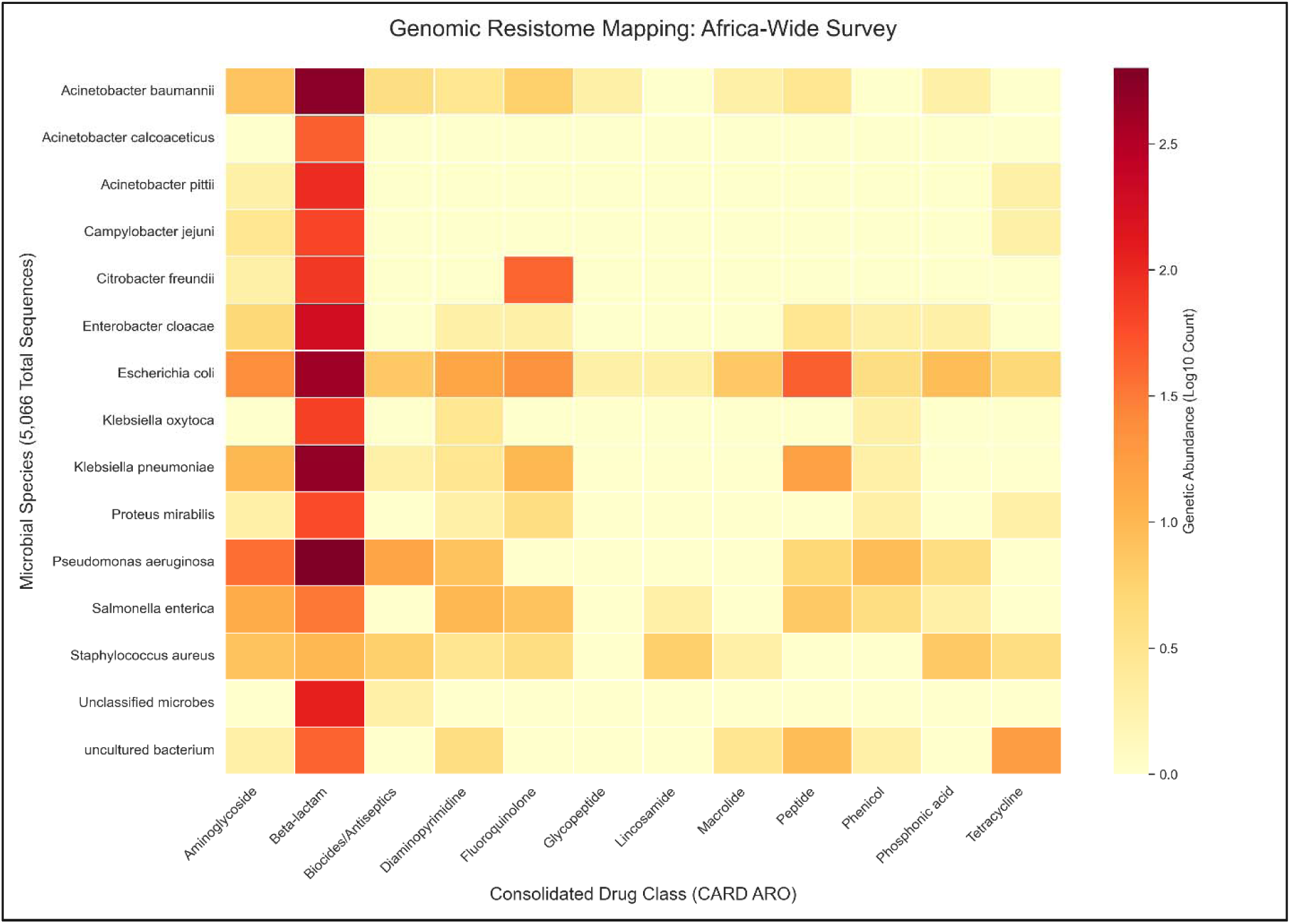
Taxonomic distribution and functional profiles of the African resistome. The heatmap illustrates the primary taxonomic drivers of antimicrobial resistance (AMR) within the Afri-ARM database, ranked by logarithm of the total antimicrobial resistance gene (ARG) counts. Color variations within each taxon represent the diversity of antibiotic drug classes.

### 3.3. African ARGs overlap with the global CARD resistome

We then sought to establish whether the African ARGs had an overlap with the global resistome in CARD. The UMAP projection of ESM-2 embeddings revealed extensive overlap between African metagenome-derived ARGs and CARD reference ARGs (Figure 3). As such, African ARGs did not form a distinct cluster or isolated region of embedding space but were interspersed throughout the global resistance landscape. In other words, African ARGs showing similar proximity to CARD ARGs as CARD ARGs show to each other. This pattern suggests that African ARGs largely occupy established functional regions associated with known resistance proteins. So, there is limited functional divergence of African ARGs from globally characterized resistance genes.

**Figure 3.**
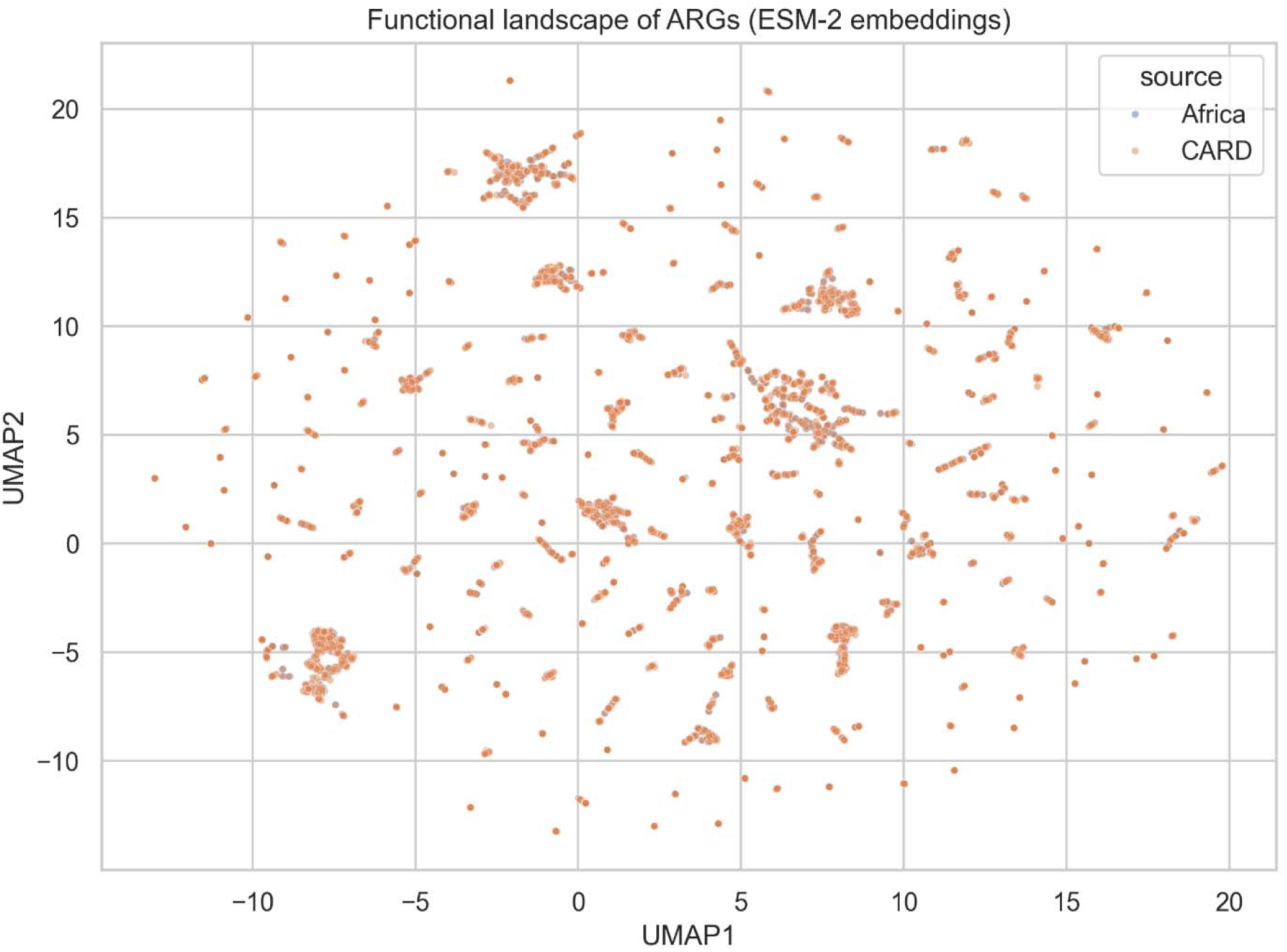
Functional landscape of antimicrobial resistance genes (ARGs) from Africa and the Comprehensive Antibiotic Resistance Database (CARD) visualized using ESM-2 protein embeddings projected by UMAP. Each point represents an ARG sequence positioned along UMAP1 (x-axis) and UMAP2 (y-axis), with coordinate ranges from −10 to 20, illustrating the shared functional embedding space between African and global ARG repertoires.

Functional density bias analysis revealed a largely uniform occupancy of protein functional space by African metagenome-derived ARGs relative to the global CARD resistome (Figure 4A). The ESM-2 embedding landscape of the African ARG density shows a balanced representation of African and CARD sequences within most regions of functional space. This pattern suggests that African ARGs do not preferentially cluster in distinct or isolated functional subspaces. Instead, they co-localize extensively with globally characterized resistance determinants. At the periphery of the embedding space, some elevated CARD-enriched density ratios were observed. However, these regions were rare and spatially restricted. Although African ARGs did not occupy a distinct functional subspace, density-based analyses also revealed non-uniform occupancy within the global resistance landscape (Figure 4B). However, these Africa-enriched regions remained embedded within the broader CARD-defined functional manifold rather than forming novel clusters. In general, the functional density landscape supports the conclusion that African ARGs largely occupy well-established resistance protein architectures with known resistance mechanisms rather than novel functional classes.

**Figure 4.**
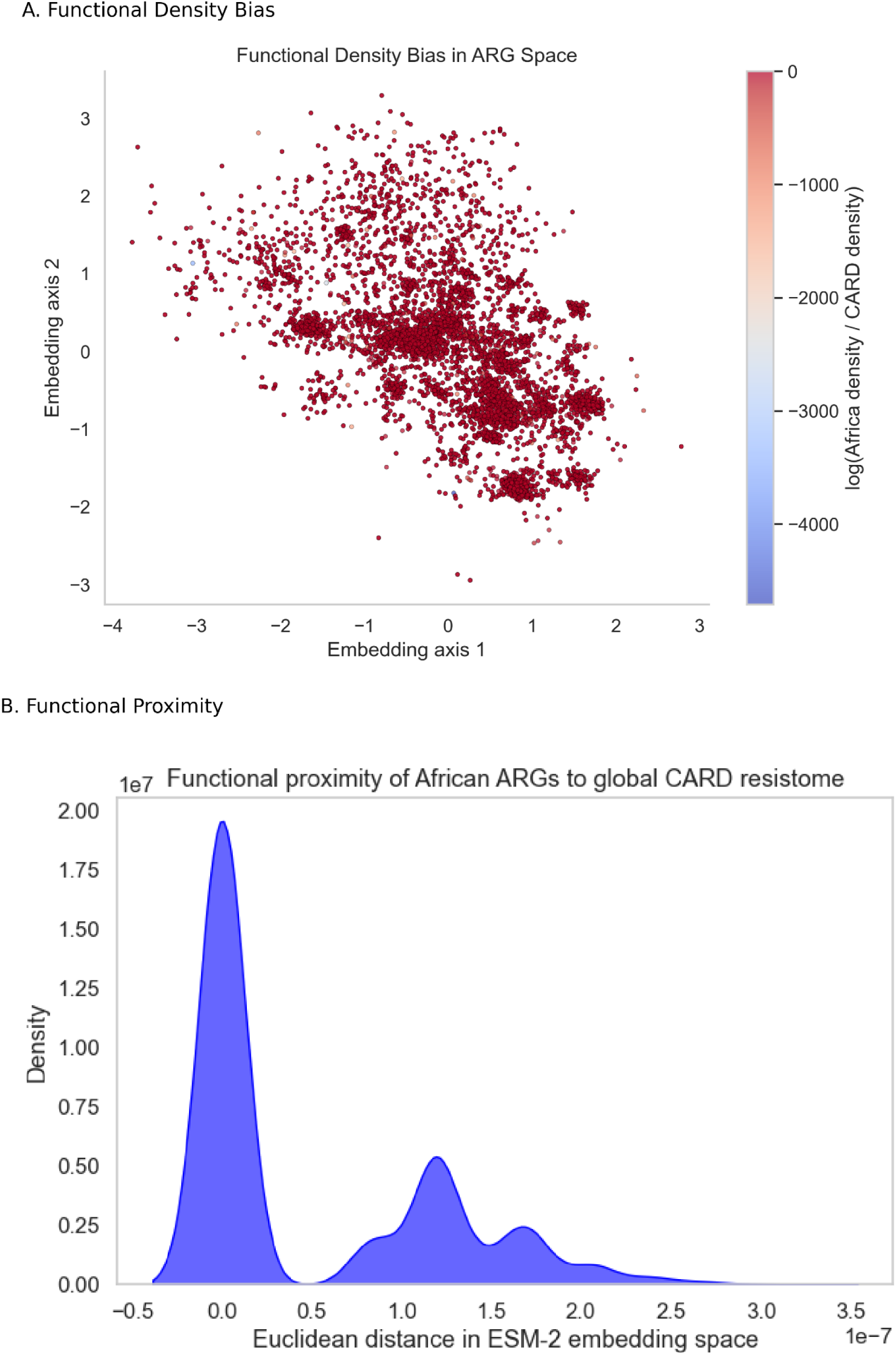
Functional bias and proximity patterns of antimicrobial resistance genes. The top panel shows the functional density bias of antimicrobial resistance genes across annotated functional categories. The bottom panel illustrates functional proximity patterns, highlighting co-occurrence relationships among resistance-associated functions.

### 3.4. Protein structure variation predicted by machine learning

The machine learning classifier showed strong discriminatory performance in distinguishing ARGs from non-ARG protein sequences based on protein-derived features. The model achieved an overall accuracy of 94.3%, which was indicative of a high proportion of correctly classified sequences across both classes. Equally, high precision (0.91) and recall (0.99) were observed for the ARG class (Figure 5). This also points to the model’s ability to reliably identify resistance-associated proteins while minimizing false negatives. The high recall value is evident that nearly all true ARGs were correctly detected. The corresponding precision reflects limited misclassification of non-ARG proteins as ARGs. The classifier also achieved an ROC AUC of 0.99. This means that the model had an excellent separability between ARG and non-ARG sequences across decision thresholds. These results indicate that protein-level features capture strong discriminatory signals associated with antimicrobial resistance. However, while the model is good at separating ARG-like proteins from curated non-ARGs, it is important to note that this does not imply prediction of novel resistance mechanisms. In this work, our model points to a classification claim and not a discovery claim.

**Figure 5.**
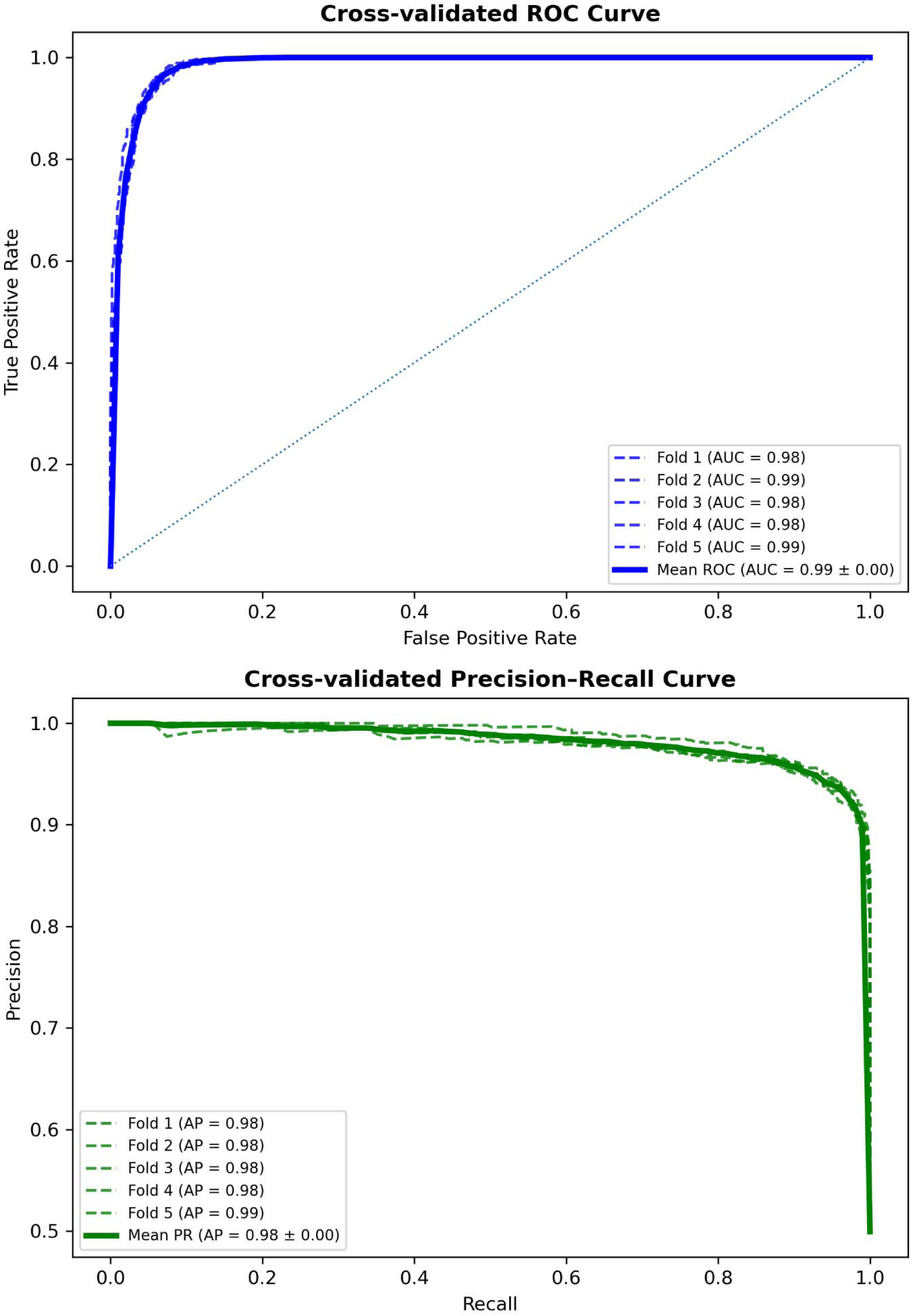
Cross-validated performance of the machine-learning model for ARG classification. The receiver operating characteristic (ROC) curve (top) summarizes the trade-off between true positive and false positive rates across cross-validation folds. The precision–recall (PR) curve (bottom) highlights the model’s ability to maintain high precision across a wide range of recall values. Shaded regions represent variability across folds, demonstrating consistent performance and minimal overfitting.

Analysis of feature importance revealed that protein compositional and physicochemical properties contributed substantially to ARG discrimination. The most important features were the fraction of tryptophan residues (AA_W; importance = 0.147), molecular weight (0.105) and protein length (0.095). This suggests that ARGs tend to occupy a distinct region of size and amino acid composition space relative to non-ARG proteins. Other influential features included the fractions of serine (AA_S) and glutamate (AA_E) residues as well as isoelectric point (pI). These features further point to the role of differences in residue polarity, charge distribution, and electrostatic properties in discriminating between ARG and non-ARG proteins. Measures related to secondary structure propensity (e.g., β-sheet fraction) and hydrophobicity (GRAVY score) further contributed to classification performance. While less prominent, their role suggests the importance of functional constraints associated with resistance mechanisms such as enzymatic antibiotic inactivation and target protection. Statistically significant differences were observed between ARG and non-ARG proteins for these top-ranked features (Mann–Whitney U test, p < 0.001). In particular, ARGs exhibited distinct distributions of tryptophan, serine, and glutamate content as well as systematically shifted isoelectric points, protein sizes, and molecular weight (Figure 6). These results indicate that ARGs are characterized by reproducible compositional and physicochemical signatures that extend beyond simple sequence similarity. Hence, it may be possible to effectively discriminate them using protein-level features.

**Figure 6.**
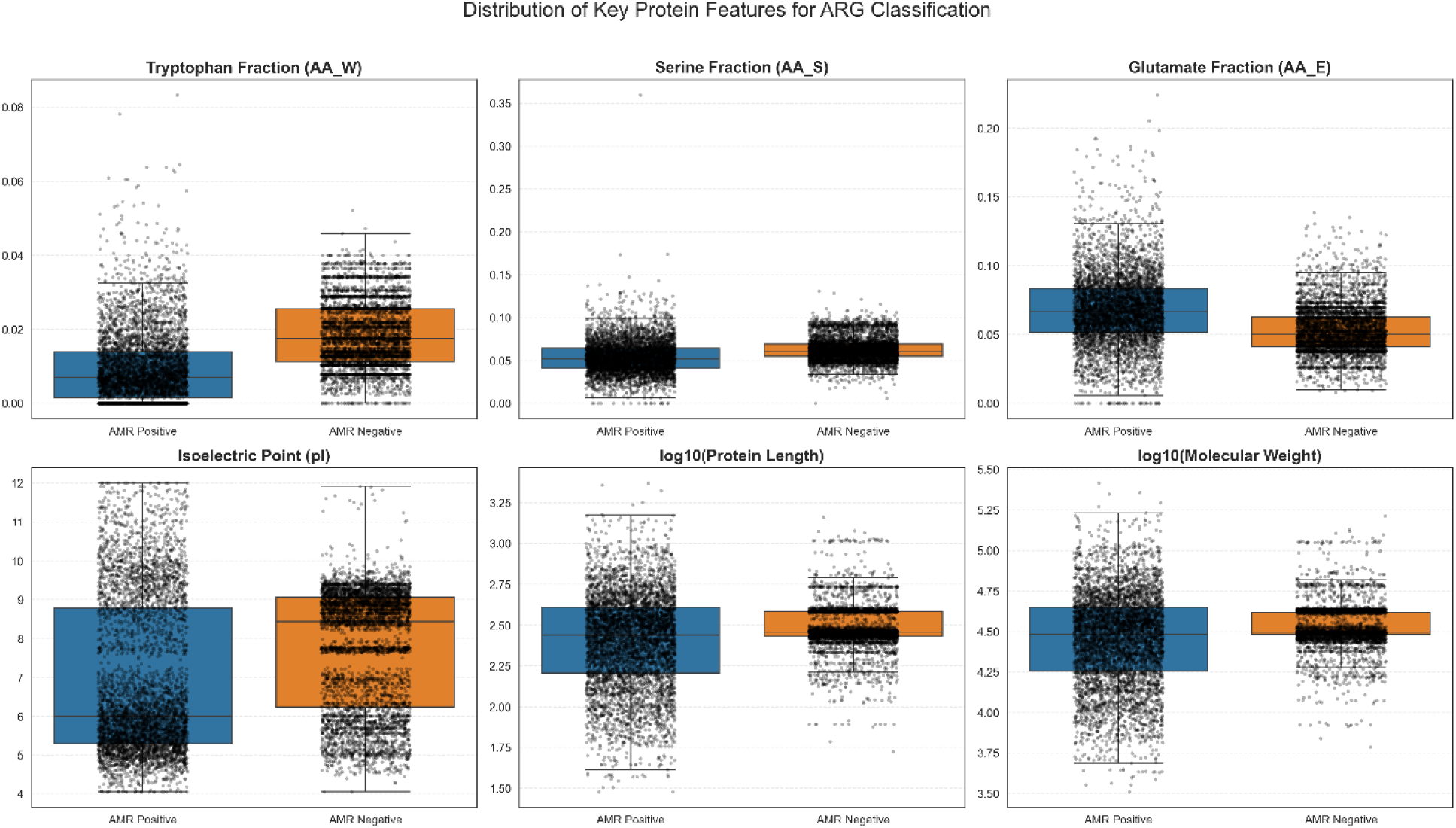
Distribution of key protein features distinguishing ARGs and non-ARGs. Box plots show the distribution of six top-ranked features—tryptophan fraction (AA_W), serine fraction (AA_S), glutamate fraction (AA_E), isoelectric point (pI), log-transformed protein length, and log10-transformed molecular weight—across ARG and non-ARG classes. Significant differences between groups were observed for all features (p < 0.001).

## 4. Discussion

This study presents a curated and harmonized African antimicrobial resistance protein resource and, in doing so, offers several insights into the structure and composition of the African resistome. A striking feature of the dataset is the predominance of ARGs associated with clinically important taxa, particularly ESKAPE pathogens such as *Pseudomonas aeruginosa*, *Acinetobacter baumannii*, and *Klebsiella pneumoniae*. This mirrors global AMR trends where ESKAPE pathogens dominate resistome burden (Ahmed et al., 2024; Ayobami et al., 2022). The trends point to the likelihood that resistance in Africa is driven largely by pathogens already entrenched in healthcare settings worldwide. Meanwhile, the overwhelming dominance of β-lactam resistance (accounting for nearly 80% of all ARG assignments) likely reflects both extensive β-lactam usage and the evolutionary plasticity of β-lactamase families. This means that African resistomes, at least at the protein level, are shaped less by unique resistance innovations and more by the expansion and diversification of globally circulating resistance mechanisms. Country-level heterogeneity, with Burkina Faso, Malawi, and Benin showing higher ARG hits per sample, further suggests that surveillance intensity, antibiotic stewardship, and sampling design may strongly influence observed resistome profiles. We postulate that these geographic differences reflect a combination of antimicrobial access patterns and uneven sequencing efforts rather than true biological signals alone.

Beyond taxonomic and functional composition, our protein embedding analysis provides an important functional perspective. By projecting African ARGs into a shared embedding space with CARD reference sequences, we observed substantial overlap between African and global resistance proteins. In practical terms, this suggests that African ARGs largely occupy the same functional subspace as previously characterized resistance determinants, arguing against the presence of widespread, structurally novel ARG classes within the analyzed datasets. This observation aligns with recent work showing that protein language models capture conserved biochemical and evolutionary constraints across diverse resistance genes (Lin et al., 2023; Rives et al., 2021). However, this interpretation should be taken cautiously. Our results are limited by the reliance on available public data and CARD-based annotation. Consequently, rare or context-specific ARGs may remain undetected, particularly those embedded in low-abundance taxa or under-sampled environments. Moreover, we do not attempt *de novo* functional validation, and therefore cannot exclude the possibility of subtle functional novelty not captured by embedding proximity alone. Nevertheless, our results support the view that resistance emergence in Africa is predominantly driven by dissemination and recombination of existing ARG families rather than the evolution of fundamentally new resistance architectures.

Finally, the machine learning framework adopted in this study highlights consistent and biologically interpretable protein-level signatures that distinguish ARGs from non-ARGs with high accuracy. The model achieved strong performance across multiple metrics, including a ROC-AUC approaching 0.99. This shows robust separability between classes. Feature importance analysis revealed that amino acid composition, protein length, molecular weight, and isoelectric point were among the most informative predictors. Further, these features differ significantly between ARGs and non-ARGs (Mann–Whitney U test, p < 0.001), which is an indication of their discriminatory value. These findings echo earlier studies showing that resistance proteins tend to occupy distinct physicochemical regimes (Arango-Argoty et al., 2018; Bhangu et al., 2025). This means that AMR patterns could be shaped by enzyme activity, substrate binding, and cellular localization. At the same time, our approach remains intentionally conservative: we do not incorporate genomic context, gene expression, or phenotypic resistance data, all of which are critical for clinical interpretation. Future work should therefore integrate multi-omic and quantitative resistance measurements to move beyond binary classification. Nonetheless, our results demonstrate that even relatively simple protein features, when combined with curated databases and modern machine learning, can yield scalable and interpretable tools for resistome analysis.

## 5. Conclusions

In this study, we examined metagenomic datasets from diverse sources in Africa. Our findings show that African metagenome-derived ARGs largely reflect globally conserved resistance functions rather than occupying distinct functional protein space. Resistance is strongly dominated by enzymatic inactivation mechanisms targeting β-lactam antibiotics. These findings suggest that AMR in African environments is shaped by ecological and usage-driven selection pressures operating within a shared global resistome. Protein language model embeddings further reinforce the view of the occupation of the same functional space with the global AMR resistome. Still, PLM offer a powerful tool for future resistome surveillance and comparative functional analyses. Using machine learning, we further show the predictive role of protein sequence structures in discriminating ARGs and non-ARGs in African metagenomes. This approach may therefore be used for future exploration of novel antimicrobial resistance genes in future surveillance efforts.

## Data Availability

All sequencing accessions analyzed in this study are publicly available from their respective ENA and SRA repositories. Processed antimicrobial resistance gene (ARG) hit data are available for download at https://www.bio-africa.org/downloads. All scripts used for data processing, feature extraction, embedding generation, and machine-learning analyses are openly available at https://github.com/ngangao/AfriARM.

## Author Contributions

**MNM**: conceptualization; formal analysis; investigation; methodology; project administration; software; validation; writing—original draft; writing—review & editing. **GWK**: visualization; writing—review & editing.

## Acknowledgements

The authors are grateful to Bio One Scientific Ltd. for the support

